# Decoding the Tumor-Suppressive Landscape of SCARA5: A Network-Based Analysis Linking Lipid Metabolism and Immune Regulation in Breast Cancer

**DOI:** 10.1101/2025.09.02.672764

**Authors:** Tooba Jawwad, Sameer Mirza, Sadaf Khursheed Baba, Harsh Kumar, Mohit Mazumder

**Affiliations:** Siriraj Center of Research Excellence for Systems Pharmacology, Department of Pharmacology, Faculty of Medicine, Siriraj Hospital, Mahidol University, Bangkok, Thailand; Department of Chemistry, College of Science (COS), United Arab Emirates University (UAEU), P.O. Box 15551, Al Ain, United Arab Emirates; OmicsLogic Inc. 10301 Northwest Freeway Ste 314 Houston, TX 77092 USA; OmicsLogic Bioinformatics & Data Science India Private Limited, #6 CGHS Plot no 7, Sec-9 Dwarka, West Delhi -110075, India

**Author notes:** Corresponding Author (T.J.).

**Keywords:** Scara5, Breast cancer subtype stratification, tumor-stroma cross-talk, single-cell and spatial transcriptomics, integrative multi-omics analysis

## Abstract

**Background:** The tumor microenvironment (TME) significantly impacts breast cancer progression, with stromal and immune components influencing tumor behavior. The scavenger receptor SCARA5 is recognized as a tumor suppressor in various cancers, but its role in breast cancer remains uncertain.

**Methods:** We performed integrative transcriptomic analyses of bulk RNA-seq data from TCGA-BRCA, validated with GTEx, to investigate SCARA5 expression and its clinical significance. Differential expression, PAM50 subtype classification, PCA/t-SNE clustering, and ROC analyses were used to assess diagnostic potential. Functional enrichment (GO, KEGG, Reactome), PPI networks, and co-expression analyses explored pathways related to SCARA5. Single-cell transcriptomic datasets (TISCH2, Broad Portal) and spatial profiling (Human Protein Atlas) were employed to examine cellular and spatial localization. The prognostic importance was assessed using GEPIA2 survival analysis.

**Results:** SCARA5 was significantly downregulated in breast tumors, especially in Her2-enriched and Luminal B subtypes, with ROC curves confirming its diagnostic importance. Enrichment and PPI analyses linked SCARA5 to lipid metabolism, immune regulation, and scavenger receptor pathways. Co-expression studies showed associations with lipid metabolism genes (FABP4, ADIPOQ, CD36) and immune-related genes (CLEC3B, LYVE1). Single-cell data indicated SCARA5 expression was limited to fibroblasts, endothelial cells, and immune subsets, with rare expression in malignant epithelial cells. Spatial analysis confirmed stromal enrichment, mainly in areas rich in fibroblasts and endothelial cells. Survival analysis demonstrated worse outcomes in patients with HER2+ and Luminal B breast cancers who had low SCARA5 expression.

**Conclusion:** SCARA5 is a stromal-enriched gene with potential tumor-suppressive and immunometabolic regulatory roles in breast cancer. Its diagnostic and prognostic significance, especially in aggressive subtypes, highlights its potential as a biomarker and therapeutic target within the tumor stroma.

## 1. INTRODUCTION

Breast cancer is the most frequently diagnosed malignancy among women worldwide, resulting in approximately 2.3 million new cases and 685,000 deaths each year (1). Despite significant advances in early detection, molecular profiling, and targeted treatments, breast cancer remains a leading cause of cancer-related death (2, 3), mainly due to its molecular diversity (4, 5) and subtype-specific severity (6). Although hormone receptor–positive subtypes typically have favorable outcomes, aggressive forms such as HER2-enriched and triple-negative breast cancer (TNBC) are linked to poor prognosis and limited treatment options (7, 8).

Recent advances have underscored the vital role of the tumor microenvironment (TME) in affecting breast cancer progression, metastasis, and treatment resistance (9, 10). The TME consists of a complex network of stromal fibroblasts, endothelial cells, immune cells, and extracellular matrix components that collectively influence tumor behavior (9, 11). Cancer-associated fibroblasts (CAFs), in particular, have been linked to promoting tumor growth, immune evasion, and metabolic reprogramming (12). Similarly, immune components within the TME can play dual roles, either supporting or inhibiting tumor progression depending on specific interactions (9, 13).

The Scavenger Receptor Class A Member 5 (SCARA5), a ferritin receptor and member of the scavenger receptor family, has been identified as a tumor suppressor in various cancers, including hepatocellular carcinoma, renal cell carcinoma, and lung cancer (14–17). SCARA5 plays a crucial role in iron homeostasis, ferritin uptake, and lipid metabolism (18–21). Recent studies have connected SCARA5 downregulation to increased tumor growth, higher metastatic potential, and immune evasion in multiple cancer types (21–23). However, despite these findings, the expression patterns, functional relationships, and clinical importance of SCARA5 in breast cancer are still not well understood. Previous research on breast cancer has linked SCARA5 downregulation to promoter hypermethylation (24) and suggested its tumor-suppressive functions may inhibit metastasis via pathways like ERK1/2, STAT3, and AKT signaling (25), but these studies were limited to small patient groups or lab models. They did not provide comprehensive analysis of SCARA5’s expression across different molecular subtypes, its prognostic value in large clinical datasets, or its role within the tumor microenvironment. Moreover, no research has systematically examined SCARA5 using integrated multi-omics approaches—combining bulk RNA-seq, single-cell transcriptomics, and spatial profiling—to clarify its stromal-specific expression and functional networks in breast cancer. In particular, its subtype-specific roles, especially in aggressive subtypes such as HER2-positive and Luminal B, remain unstudied. Additionally, there is a lack of systems biology analyses to understand its regulatory context, including possible links to pathways like PPAR signaling. The spatial distribution and single-cell heterogeneity of SCARA5 within the breast tumor microenvironment (TME) also remain largely unknown, representing a significant knowledge gap.

With the rise of high-throughput omics technologies, integrative transcriptomic analyses that combine bulk RNA sequencing (RNA-seq), single-cell RNA-seq (scRNA-seq), and spatial transcriptomics have enabled a deeper understanding of gene regulation within the TME (26–31). Tools like PAM50 molecular subtyping, functional enrichment analyses, and protein–protein interaction (PPI) networks further help identify gene signatures with potential diagnostic and prognostic value (32, 33).

In this study, we used a multi-omics approach, utilizing TCGA-BRCA bulk RNA-seq data, single-cell transcriptomic datasets from TISCH2 and the Broad Institute Single Cell Portal, and spatial expression profiles from the Human Protein Atlas to systematically examine the expression landscape of SCARA5 in breast cancer. We also investigated its connection to lipid metabolism, immune regulatory pathways, and patient survival outcomes. Our results show that SCARA5 is mainly expressed in stromal fibroblasts and endothelial cells, indicating a potential role in tumor-stroma interactions and emphasizing its importance as a prognostic marker in aggressive breast cancer subtypes.

## 2. MATERIALS and METHODS

### 2.1 Data Collection and Sources

An integrated systems biology approach was employed to investigate the expression profile, prognostic significance, co-expression network, and spatial localization of SCARA5 in breast cancer, utilizing publicly available databases and computational tools. Transcriptomic data for breast cancer and corresponding normal tissues were obtained from The Cancer Genome Atlas (TCGA-BRCA) (https://portal.gdc.cancer.gov/projects/TCGA-BRCA) and the Genotype Tissue Expression (GTEx) project through the UCSC Xena platform (https://xenabrowser.net/datapages/?cohort=GTEX). Subtype-specific analyses were conducted based on PAM50 classification. For cell-type-specific gene expression at single-cell resolution, the Tumor Immune Single-cell Hub (TISCH2) (http://tisch.comp-genomics.org/home/) was employed. Single-cell RNA sequencing analyses were performed using the Single Cell Portal (https://singlecell.broadinstitute.org/single_cell) (TISCH). Survival analysis was conducted with GEPIA2 (http://gepia2.cancer-pku.cn/#index). Immunohistochemical and spatial transcriptomics data were sourced from the Human Protein Atlas (HPA) (https://www.proteinatlas.org/ENSG00000168079-SCARA5/single+cell/breast). Differential gene expression analysis, SCARA5 expression in tumor versus normal tissue, ROC curve analysis, pathway analysis, and protein–protein interaction studies were performed using the R programming language.

### 2.2 Expression Analysis

Differential expression analysis was conducted between tumor and normal samples using the DESeq2 package in R, based on FPKM files downloaded from the TCGA-BRCA cohort (GDC Data Portal). Normal breast tissue expression data from the GTEx database were used for validation. The differentially expressed genes were identified with a significance threshold of |log2 Fold Change| > 1 and adjusted p-value (q-value) < 0.05 (Benjamini-Hochberg correction). To validate SCARA5 mRNA expression in breast cancer, we also queried GEPIA2 (http://gepia2.cancer-pku.cn) using the "Expression DIY" module, comparing TCGA-BRCA tumor samples with GTEx normal breast tissues. Expression values were normalized as Transcripts Per Million (TPM) and log2-transformed before comparison, with a significance threshold of |log2 Fold Change| > 1 and adjusted p-value (q-value) < 0.05. For breast cancer subtype stratification, PAM50 molecular classification was applied to TCGA-BRCA RNA-seq data using a custom R script based on the published PAM50 gene set and subtype centroids. Subtypes analyzed included Luminal A, Luminal B, HER2-enriched, Basal-like, and Normal-like.

### 2.3 Dimensionality Reduction and Clustering Analysis

To investigate the distribution of breast cancer subtypes based on gene expression profiles, we used Principal Component Analysis (PCA) and t-Distributed Stochastic Neighbor Embedding (t-SNE). For overall expression patterns, the top 50 differentially expressed genes (DEGs) identified from TCGA-BRCA were selected. Expression data were scaled and centered before performing PCA with the prcomp function in R, and PCA biplots were visualized using factoextra. The t-SNE analysis was conducted using the Rtsne package with a perplexity of 30 and 500 iterations, projecting the expression profiles into two dimensions to improve subtype visualization. These analyses facilitated the visualization of sample clustering patterns in relation to PAM50 molecular subtypes.

### 2.4 ROC Curve Analysis

To assess the diagnostic ability of SCARA5 in differentiating breast cancer subtypes from normal tissues, Receiver Operating Characteristic (ROC) curve analysis was carried out using the pROC and ggplot2 packages in R. The analysis was performed for each PAM50-defined subtype (Basal, HER2-enriched, Luminal A, and Luminal B) versus normal breast tissues. For each comparison, tumor samples from a specific subtype and normal samples were combined, and binary class labels were assigned (Tumor = 0, Normal = 1). The Area Under the Curve (AUC) and 95% Confidence Intervals (CI) were calculated using the DeLong method. ROC curves were visualized with ggroc() from the pROC package, and AUC values were reported for each subtype. All ROC curves and AUCs were plotted together to evaluate SCARA5’s ability to distinguish between different breast cancer subtypes.

### 2.5 Pathway Enrichment and Functional Network Analysis

To explore the biological significance of the top differentially expressed genes (DEGs) related to SCARA5 expression, we conducted functional enrichment analyses using GO Biological Processes (GO-BP), KEGG pathways, and Reactome pathways. The top 50 DEGs were converted to Entrez IDs using the bitr function from the clusterProfiler package, and enrichment analyses were performed with the enrichGO, enrichKEGG, and enrichPathway functions, applying a Benjamini-Hochberg corrected p-value < 0.05 as the significance threshold. Barplots and dotplots were created to visualize enriched pathways related to lipid metabolism, adipogenesis, and immune response. To assess the expression patterns of genes in enriched pathways, we used voom-normalized expression data (limma + edgeR pipeline) for hierarchical clustering and heatmap generation with the pheatmap package. Heatmaps were produced for both Reactome-enriched genes and combined enrichment results across GO, KEGG, and Reactome datasets. Additionally, a functional interaction network was built based on literature-supported biological relationships between SCARA5 and its associated pathways, using the igraph and ggraph packages. Nodes represented genes, biological processes, pathways, or outcomes, while directed edges depicted known interactions or regulatory effects. The network highlighted the connections between SCARA5, lipid metabolism, immune modulation, and tumor suppression in breast cancer.

### 2.6 Protein–Protein Interaction (PPI) Network and Co-expression Analysis

To explore potential interactions and co-regulation among SCARA5 and its related genes, we performed a multi-level analysis combining protein–protein interaction data, gene expression profiling, and correlation analysis. The STRING database (v11.5) was used to construct a PPI network centered on SCARA5 and its known interactors or top co-expressed genes. Gene symbols were mapped to STRING IDs using the STRINGdb R package, and interaction data with a confidence score above 400 were retrieved. The network was visualized using igraph, ggraph, and tidygraph packages, with node attributes indicating functional categories such as metabolism, immune response, extracellular matrix (ECM) remodeling, and stress response. For co-expression analysis, the expression matrix of SCARA5 and its interactors was extracted from the voom-normalized data. Heatmaps showing gene expression patterns across tumor and normal samples were created with the pheatmap package, applying hierarchical clustering to both genes and samples. Pearson correlation analysis between SCARA5 and its interactors was performed on the expression matrix, and the resulting correlation matrix was visualized as a clustered heatmap with correlation coefficients displayed. This comprehensive approach helped us map the interaction landscape of SCARA5, highlighting its association with metabolic, immune, and ECM-related gene networks in breast cancer.

### 2.7 Survival Analysis

Overall survival (OS) analysis for SCARA5 expression in breast cancer was performed using the GEPIA2 online tool (http://gepia2.cancer-pku.cn). Patients were divided into two groups based on the median expression threshold: high and low expression groups. Kaplan–Meier survival curves were generated, and statistical significance was evaluated using the log-rank test. Hazard ratios (HR) with 95% confidence intervals and corresponding p-values were obtained directly from the GEPIA2 survival module. Subtype-specific survival analysis was also conducted for the Basal-like, HER2-enriched, Luminal A, and Luminal B subtypes using the GEPIA2 “Survival Analysis” module with PAM50 subtype filters applied.

### 2.8 Single-cell Transcriptomic and Spatial Heterogeneity Analysis of SCARA5 Expression

To investigate the cell–type–specific expression patterns and spatial heterogeneity of SCARA5 in breast cancer, we performed a comprehensive single-cell transcriptomic analysis using two publicly available resources: TISCH2 (http://tisch.comp-genomics.org/) and the Broad Institute Single Cell Portal (https://singlecell.broadinstitute.org/single_cell). In TISCH2, we explored annotated single-cell RNA-seq datasets, including BRCA_Alex, BRCA_GSE114727_inDrop, and BRCA_GSE176078, which provided curated cell-type classifications within the breast cancer microenvironment. We assessed SCARA5 expression across major cellular compartments—immune cells, stromal cells, and malignant epithelial cells—using UMAP projections, violin plots, and cell composition barplots generated directly from the platform interface. Additionally, the Broad Institute Single Cell Portal was used to validate SCARA5 expression patterns across independent datasets, focusing on cancer epithelial cells, cancer-associated fibroblasts (CAFs), endothelial cells, and immune cell populations. Gene expression distributions were visualized using UMAP plots, violin plots, and dot plots integrated within the portal.

To further evaluate the spatial heterogeneity of SCARA5 expression across breast tissue cell types, we utilized data from the Human Protein Atlas (HPA) (https://www.proteinatlas.org/). This analysis included breast-specific cell populations such as glandular epithelial cells, progenitor cells, myoepithelial cells, fibroblasts, endothelial cells, adipocytes, and immune cells. We examined cell-type enrichment plots and gene correlation heatmaps provided by the HPA to determine the specificity and co-expression of SCARA5 with lineage-defining markers.

All analyses were performed using R (v4.2.0) and web-accessible tools. All data used are publicly available and accessible, ensuring full reproducibility.

## 3. RESULTS

### 3.1 SCARA5 is Significantly Downregulated in Breast Cancer Compared to Normal Tissues

We performed differential expression analysis on the TCGA-BRCA cohort, using DESeq2, to compare tumor samples with adjacent normal tissues. Among the significantly dysregulated genes (|log2 Fold Change| > 1; q-value < 0.05), SCARA5 emerged as one of the top downregulated candidates (Figure 1B). The volcano plot displayed a significant negative log2 fold change for SCARA5, suggesting its marked suppression in breast tumors.

**Figure 1.**
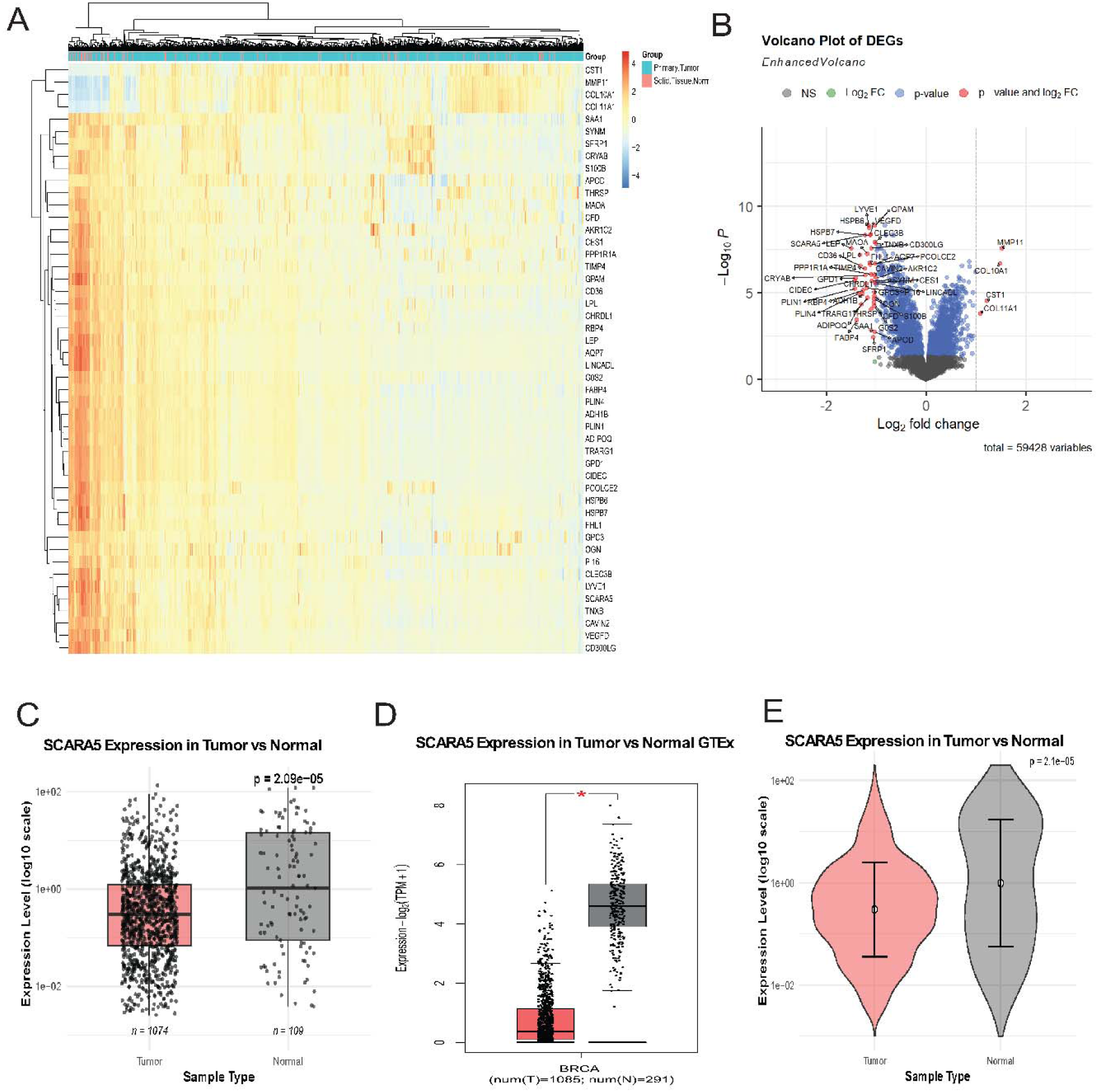
SCARA5 is significantly downregulated in breast cancer tissues. **(A)** Hierarchical clustering heatmap of the top 50 differentially expressed genes (DEGs) in the TCGA-BRCA cohort showing distinct segregation of tumor and normal samples, with SCARA5 clustering among consistently downregulated genes. **(B)** Volcano plot illustrating DEGs with SCARA5 highlighted as a significantly downregulated gene. **(C–D)** Boxplots depicting SCARA5 expression levels in tumor versus normal tissues in the TCGA-BRCA dataset (**C**) and GTEx normal breast tissues (**D**), confirming significant downregulation in tumors (p < 0.005). **(E)** Violin plot comparing SCARA5 expression between tumor and normal tissues, demonstrating a consistent reduction in tumors.

Hierarchical clustering analysis of the top 50 differentially expressed genes (DEGs) further confirmed a distinct separation between tumor and normal samples, with SCARA5 clustering among the genes consistently downregulated in tumors (Figure 1A). The full list of the top 50 differentially expressed genes used in clustering and further analyses is provided in Supplementary Table S1.

To validate these findings, we queried the GEPIA2 web server, which integrates data from TCGA and GTEx. Consistent with our in-house analysis, SCARA5 expression was significantly lower in breast cancer tissues compared to both TCGA normal (Figure 1C) and GTEx-derived normal breast tissues (Figure 1D), with p-values < 0.05. Violin plot comparisons reinforced these observations, revealing distinct expression distributions between tumor and normal samples (Figure 1E).

Collectively, these results confirm that SCARA5 is significantly downregulated in breast tumors and may contribute to a gene signature distinguishing malignant from normal breast tissues.

### 3.2 SCARA5 is Consistently Downregulated Across Molecular Subtypes of Breast Cancer

To determine whether SCARA5 downregulation is subtype-specific, we stratified the TCGA-BRCA cohort using PAM50 molecular classification, categorizing tumors into Basal-like, HER2-enriched, Luminal A, and Luminal B subtypes (Figure 2A).

**Figure 2.**
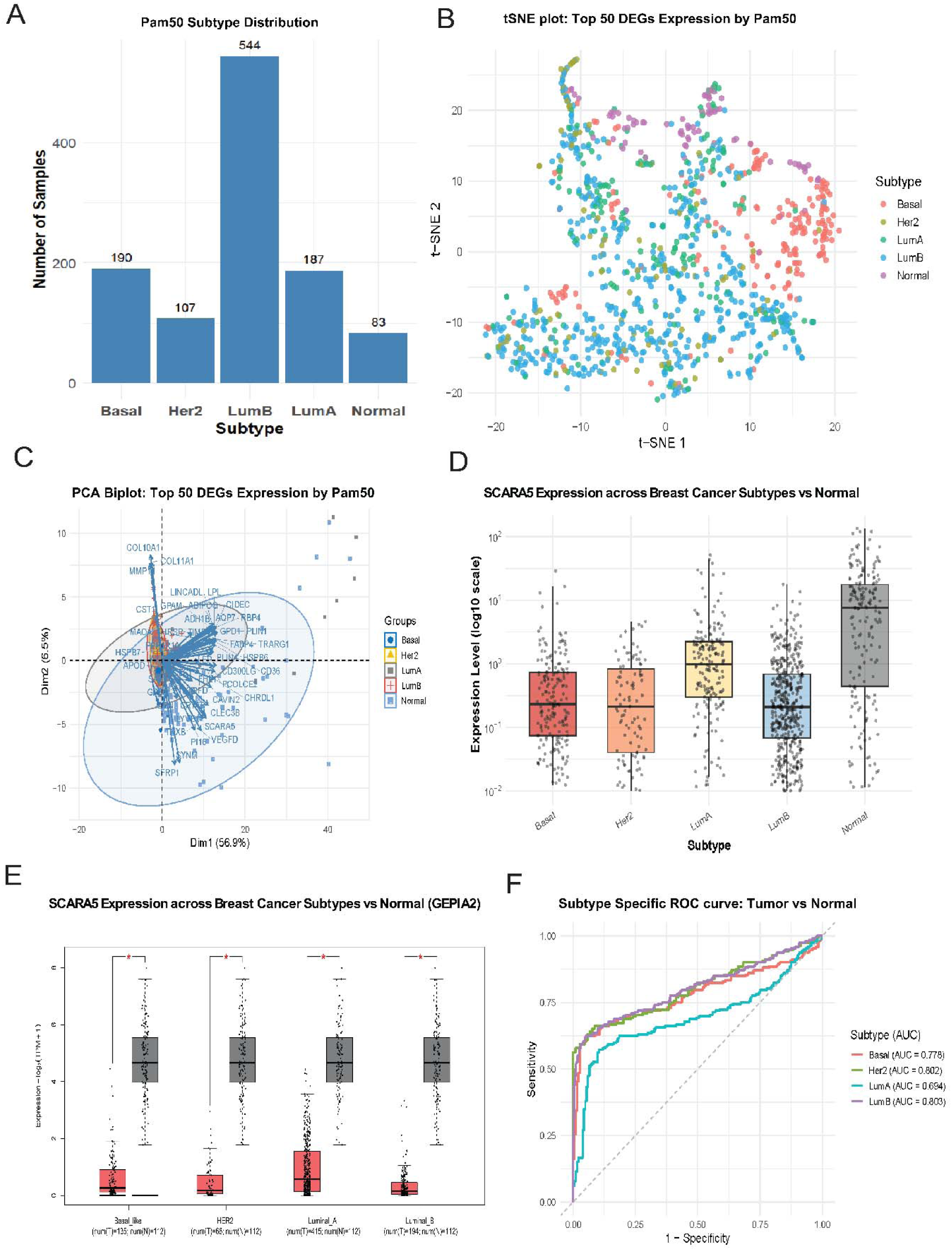
SCARA5 expression is consistently reduced across breast cancer subtypes and demonstrates diagnostic potential. (A) Bar plot showing PAM50 subtype distribution of breast cancer and normal samples. (B) t-Distributed Stochastic Neighbor Embedding (t-SNE) plot based on the top 50 differentially expressed genes (DEGs), demonstrating subtype-specific clustering. (C) Principal Component Analysis (PCA) biplot using the top 50 DEGs, showing distinct separation between PAM50-defined subtypes and normal tissues. (D) Boxplot of SCARA5 expression across PAM50 breast cancer subtypes and normal tissues, showing significant downregulation in HER2-enriched and Luminal B tumors. (E) Independent validation of SCARA5 downregulation using GEPIA2 expression data across PAM50 subtypes. (F) Subtype-specific receiver operating characteristic (ROC) curves showing the diagnostic performance of SCARA5 in distinguishing breast cancer from normal tissues (AUC range: 0.778–0.803).

Expression analysis revealed that SCARA5 expression was consistently decreased across all breast cancer subtypes compared to normal samples, with the most pronounced reduction observed in HER2-enriched and Luminal B tumors (Figure 2D). Validation with GEPIA2 showed similar patterns of reduced SCARA5 expression across subtypes (Figure 2E).

To explore subtype-specific transcriptional patterns, we applied t-distributed Stochastic Neighbor Embedding (t-SNE) and Principal Component Analysis (PCA) on the top 50 DEGs. Both dimensionality reduction methods clearly segregated tumor samples according to PAM50 subtypes, with SCARA5 contributing to subtype-specific clustering patterns (Figure 2B–C). These analyses highlighted the ability of SCARA5 to reflect underlying molecular heterogeneity within breast cancer subtypes.

### 3.3 SCARA5 Shows Robust Diagnostic Potential Across Breast Cancer Subtypes

To assess the diagnostic potential of SCARA5, we performed Receiver Operating Characteristic (ROC) curve analysis comparing each PAM50-defined subtype against normal breast tissues. The analysis showed Area Under the Curve (AUC) values ranging from 0.778 to 0.803 across Basal-like, HER2-enriched, Luminal A, and Luminal B subtypes (Figure 2F). These values reflect a strong discriminatory ability of SCARA5 to distinguish tumors from normal tissues, particularly in HER2-enriched and Luminal B tumors, where both expression differences and AUC values were highest.

The consistent performance of SCARA5 across distinct molecular subtypes suggests its potential as a pan-subtype biomarker with possible utility in breast cancer diagnosis and subtype-specific profiling.

### 3.4 Pathway Enrichment and Functional Network Analysis Reveal SCARA5-Associated Metabolic and Immune Regulatory Roles

To elucidate the biological processes dysregulated in breast cancer and potentially linked to SCARA5 function, we performed pathway enrichment analysis on the top 50 differentially expressed genes (DEGs) identified between tumor and normal breast tissues. Using GO Biological Processes (GO-BP), KEGG, and Reactome databases, we observed significant enrichment in pathways related to lipid metabolism, adipocyte differentiation, immune modulation, and scavenger receptor activity (Figure 3A–C). The detailed list of Reactome-enriched pathways is provided in Supplementary Table S2.

**Figure 3.**
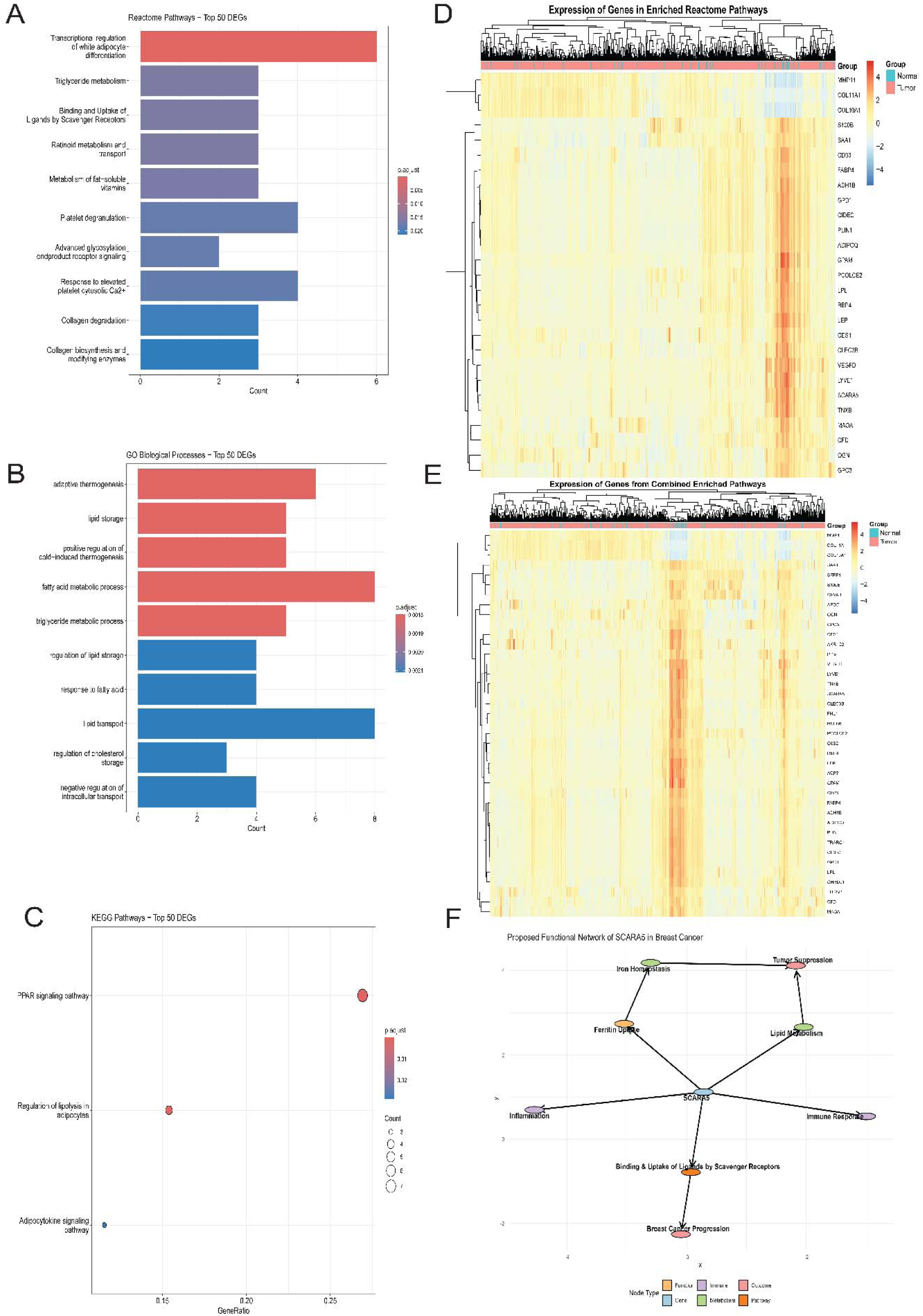
Functional enrichment analyses link SCARA5 to pathways involved in lipid metabolism and immune regulation. **(A)** Reactome pathway enrichment of the top 50 differentially expressed genes (DEGs), highlighting key biological processes including lipid metabolism, scavenger receptor activity, and immune regulation. **(B)** KEGG pathway enrichment analysis showing involvement of DEGs in PPAR signaling, adipocytokine signaling, and lipid metabolic pathways. **(C)** Gene Ontology Biological Process (GO-BP) analysis demonstrating enrichment in lipid transport, fatty acid metabolism, and immune-modulatory processes. **(D–E)** Heatmaps depicting expression profiles of enriched pathway genes, showing a consistent downregulation pattern in tumor samples compared to normal tissues. **(F)** Proposed functional interaction network integrating SCARA5 within lipid metabolism, immune regulation, iron homeostasis, and tumor suppression pathways in breast cancer.

Among the Reactome pathways, DEGs were predominantly enriched in transcriptional regulation of white adipocyte differentiation, ligand binding and uptake by scavenger receptors, and triglyceride metabolism (Figure 3A). Genes mapped to these enriched Reactome pathways are detailed in Supplementary Table S3. KEGG analysis further highlighted key metabolic pathways, including PPAR signaling, adipocytokine signaling, and regulation of lipolysis in adipocytes (Figure 3B). Consistently, GO-BP enrichment showed overrepresentation of terms such as fatty acid metabolic process, lipid transport, and lipid storage regulation (Figure 3C). The combined pathway enrichment results from GO-BP, KEGG, and Reactome analyses are summarized in Supplementary Table S4.

Expression heatmaps of genes from these enriched pathways revealed a consistent downregulation pattern in tumor samples compared to normal tissues (Figure 3D–E). Notably, SCARA5 clustered with several metabolic regulators, suggesting a coordinated repression of metabolic and immune-regulatory genes in the tumor setting.

Finally, we constructed a functional interaction network to integrate SCARA5 within these enriched biological contexts. The network positioned SCARA5 at the intersection of lipid metabolism, immune regulation, iron homeostasis, and tumor suppression, supporting a model where the repression of SCARA5 and related pathways may contribute to metabolic reprogramming and immune evasion in breast cancer (Figure 3F).

### 3.5 SCARA5 Co-expression Network Reveals Distinct Metabolic and Immune Interactors

To further explore the regulatory landscape of SCARA5, we conducted a comprehensive analysis that integrated protein–protein interaction (PPI) networks, gene expression heatmaps, and co-expression patterns. The PPI network, derived from STRING database interactions, identified two primary gene clusters associated with SCARA5: (1) Metabolism-related genes (FABP4, ADIPOQ, LEP, GPAM, CD36), known for roles in lipid metabolism and adipogenesis. (2) Immune/ECM-related genes (CLEC3B, LYVE1, TNXB), implicated in extracellular matrix remodeling and immune regulation (Figure 4A–B). Expression heatmaps of these interactors showed uniform downregulation in tumor tissues (Figure 4C), mirroring the pattern observed for SCARA5, which suggests possible co-regulation under tumor-specific transcriptional programs.

**Figure 4.**
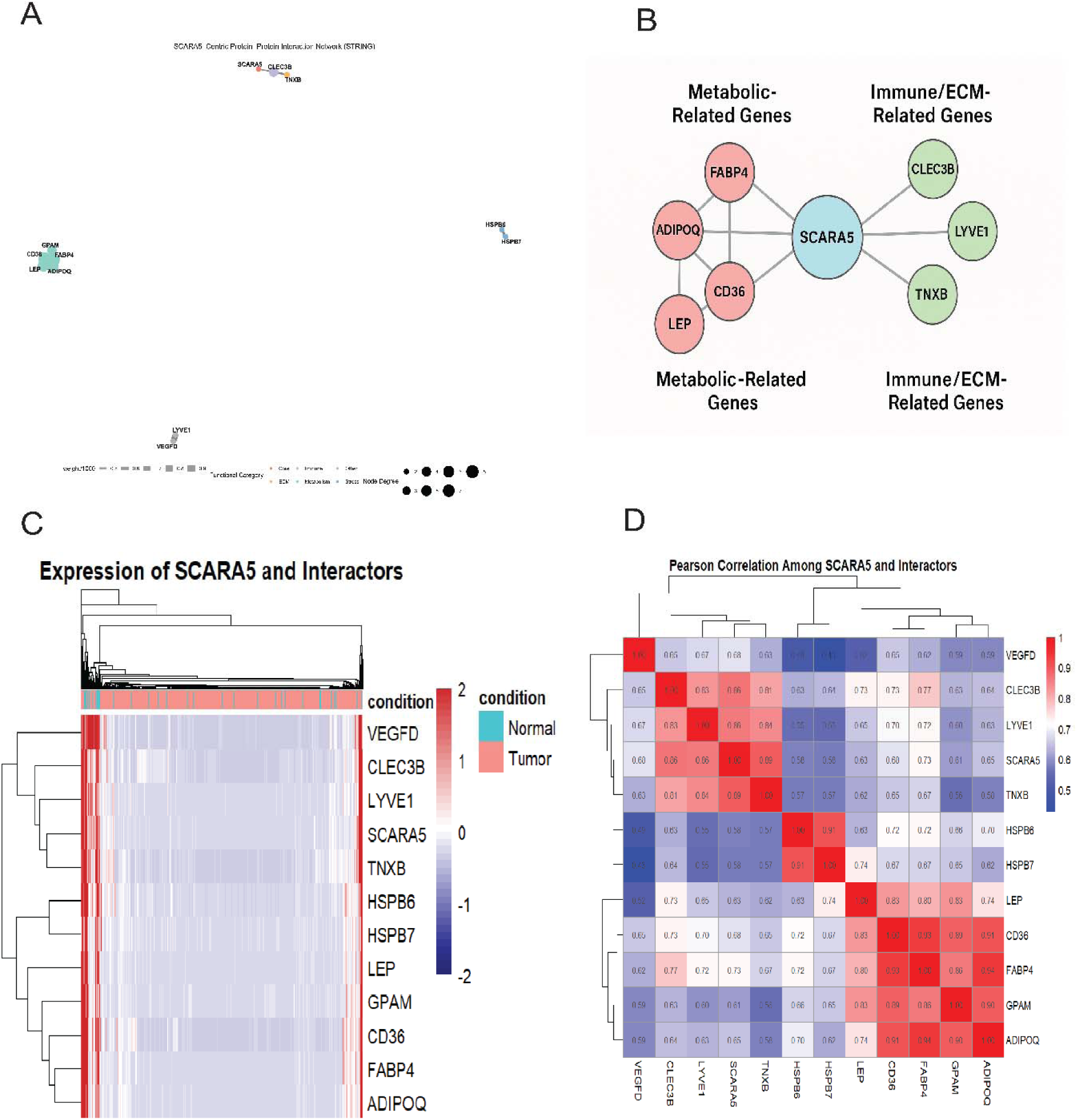
Co-expression and protein–protein interaction networks of SCARA5 reveal distinct metabolic and immune interactors. **(A)** STRING-based protein–protein interaction (PPI) network of SCARA5 and its interactors, highlighting clusters associated with metabolism and immune/ECM functions. **(B)** Subnetwork depicting metabolic interactors (e.g., FABP4, ADIPOQ, CD36) and immune/ECM interactors (e.g., CLEC3B, LYVE1, TNXB). **(C)** Heatmap showing expression profiles of SCARA5 and its interactors across tumor and normal tissues. **(D)** Pearson correlation matrix indicating strong co-expression among metabolic genes and moderate correlations with immune-related genes.

Pearson correlation analysis revealed strong positive correlations within the metabolic cluster (r > 0.9), and moderate-to-strong correlations between SCARA5 and immune/ECM-related genes (r = 0.6–0.7) (Figure 4D). This high degree of co-expression indicates that SCARA5 is embedded within tightly regulated metabolic and immune gene networks, potentially orchestrated by shared transcriptional regulators such as PPAR signaling pathways.

These findings underscore SCARA5’s integrative role at the interface of metabolism and immune response, with its downregulation in breast tumors suggesting a functional link to disrupted cellular homeostasis and tumor progression.

### 3.6 SCARA5 Expression is Linked to Subtype-Specific Prognosis in Breast Cancer

To investigate the prognostic relevance of SCARA5 expression in breast cancer, we performed overall survival (OS) analysis using the GEPIA2 survival analysis module, categorizing patients into high- and low-expression groups based on median SCARA5 expression levels. In the overall cohort, SCARA5 expression showed a non-significant trend toward poorer survival in patients with reduced expression (HR = 0.91; p = 0.55) (Figure 5A).

**Figure 5.**
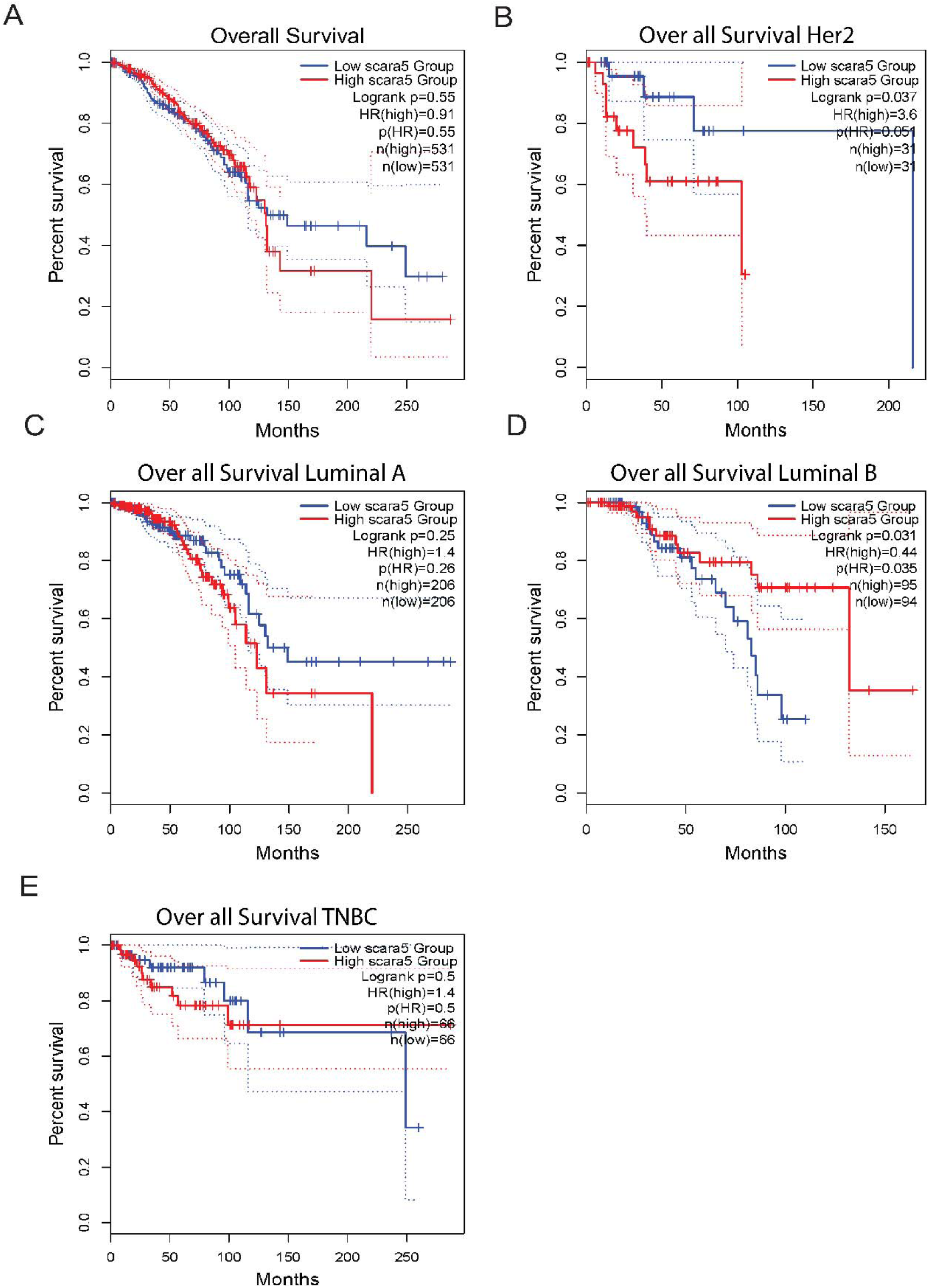
Prognostic significance of SCARA5 expression across breast cancer subtypes. **(A)** Kaplan–Meier curve showing overall survival (OS) for the TCGA-BRCA cohort based on SCARA5 expression (non-significant trend). **(B–E)** Subtype-specific OS analyses reveal significant associations between low SCARA5 expression and poor prognosis in HER2-enriched (**C**; HR = 0.34, p = 0.03) and Luminal B (**E**; HR = 0.63, p = 0.04) subtypes, with no significant impact in Basal-like (**B**) or Luminal A (**D**) subtypes.

However, when stratified by PAM50 subtypes, distinct prognostic patterns emerged. In the HER2-enriched subgroup, low SCARA5 expression was associated with markedly poorer survival outcomes (HR = 0.34; p = 0.03) (Figure 5C). Similarly, in Luminal B tumors, patients with reduced SCARA5 expression exhibited significantly worse overall survival (HR = 0.63; p = 0.04) (Figure 5E). No significant survival associations were observed in the Basal-like (HR = 0.74; p = 0.14) (Figure 5B) or Luminal A (HR = 0.84; p = 0.25) (Figure 5D) subtypes.

These findings suggest that SCARA5 exhibits subtype-specific prognostic significance, particularly in HER2-enriched and Luminal B breast cancers, where its reduced expression is associated with poorer clinical outcomes.

### 3.7 SCARA5 is Selectively Expressed in the Stromal and Immune Compartments of the Breast Tumor Microenvironment

To dissect the cellular context of SCARA5 expression in the breast tumor ecosystem, we performed a comprehensive single-cell transcriptomic analysis using curated datasets from TISCH2 and the Broad Institute Single Cell Portal. UMAP clustering of breast cancer single-cell datasets, including BRCA_Alex and BRCA_GSE114727_inDrop, revealed distinct cellular populations comprising malignant epithelial cells, stromal cells (fibroblasts, endothelial cells), and various immune subsets (T cells, macrophages, NK cells) (Figure 6A, 6D).

**Figure 6.**
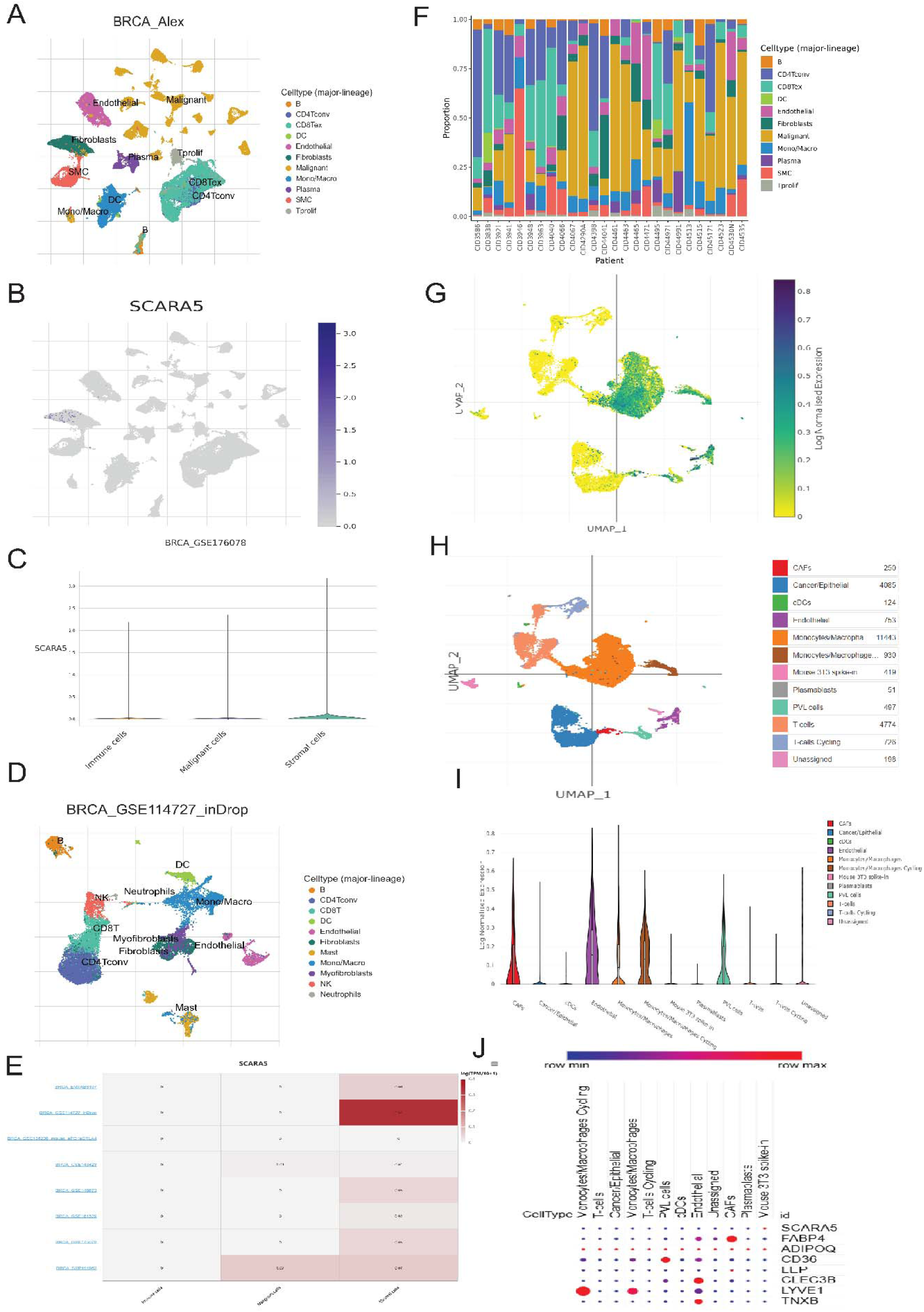
Single-cell transcriptomics highlight stromal-enriched expression of SCARA5 in the breast tumor microenvironment. **(A)** UMAP projection from single-cell dataset (TISCH2) showing cellular composition of the breast tumor microenvironment. **(B)** Violin plot quantifying SCARA5 expression levels across different cellular compartments, demonstrating stromal cell enrichment. **(C)** Feature plot mapping SCARA5 expression, revealing its enrichment in fibroblasts, endothelial cells, and select immune subsets. **(D)** UMAP projection showing detailed cellular annotations within the tumor microenvironment. **(E)** Heatmap displaying SCARA5 expression patterns across different cell types and patient samples. **(F)** Barplot depicting inter-patient variability in stromal cell abundance expressing SCARA5. **(G)** UMAP visualization colored by SCARA5 expression intensity across all cell types. **(H)** UMAP projection showing major cell type classifications with detailed annotations and cell counts. **(I)** Violin plots comparing SCARA5 expression across different cell type categories. **(J)** Dot plot illustrating co-expression of SCARA5 with lipid metabolism and immune-modulatory genes across stromal compartments.

Mapping SCARA5 expression onto these UMAP plots highlighted its near absence in malignant epithelial clusters while revealing prominent expression within fibroblasts, endothelial cells, and select immune cell populations (Figure 6B). This trend was further validated across additional datasets, including BRCA_GSE176078, where violin plots demonstrated significantly higher SCARA5 expression within stromal compartments, compared to immune and malignant epithelial compartments (Figure 6C).

Dot plot analyses from TISCH2 consistently showed SCARA5 co-expression with key lipid metabolism regulators (e.g., FABP4, ADIPOQ, GPAM, CD36) and immune-modulatory genes (CLEC3B, LYVE1, TNXB) across fibroblasts and macrophage-like cells (Figure 6J). Furthermore, proportional bar plots demonstrated patient-level heterogeneity in the abundance of SCARA5-expressing stromal cells, underscoring the variability of stromal involvement across individual tumors (Figure 6F).

Collectively, these findings indicate that SCARA5 expression is highly compartmentalized within the breast tumor microenvironment, predominantly marking stromal fibroblasts and endothelial cells, with limited expression in immune cells, and is largely absent in malignant epithelial populations. This cellular specificity suggests a non-tumor-autonomous role for SCARA5, potentially mediating stromal remodeling, immune regulation, or metabolic crosstalk within the tumor niche.

### 3.8 Spatial Heterogeneity Highlights SCARA5’s Fibroblast and Endothelial Cell Localization in Breast Tissue

To further characterize the spatial expression patterns of SCARA5 within the breast tissue landscape, we interrogated Human Protein Atlas (HPA) spatial transcriptomics datasets. Cell-type enrichment analyses revealed that fibroblasts and endothelial cells exhibited the highest SCARA5 expression scores, surpassing those of adipocytes, immune cells, and various epithelial subtypes (Figure 7A).

**Figure 7.**
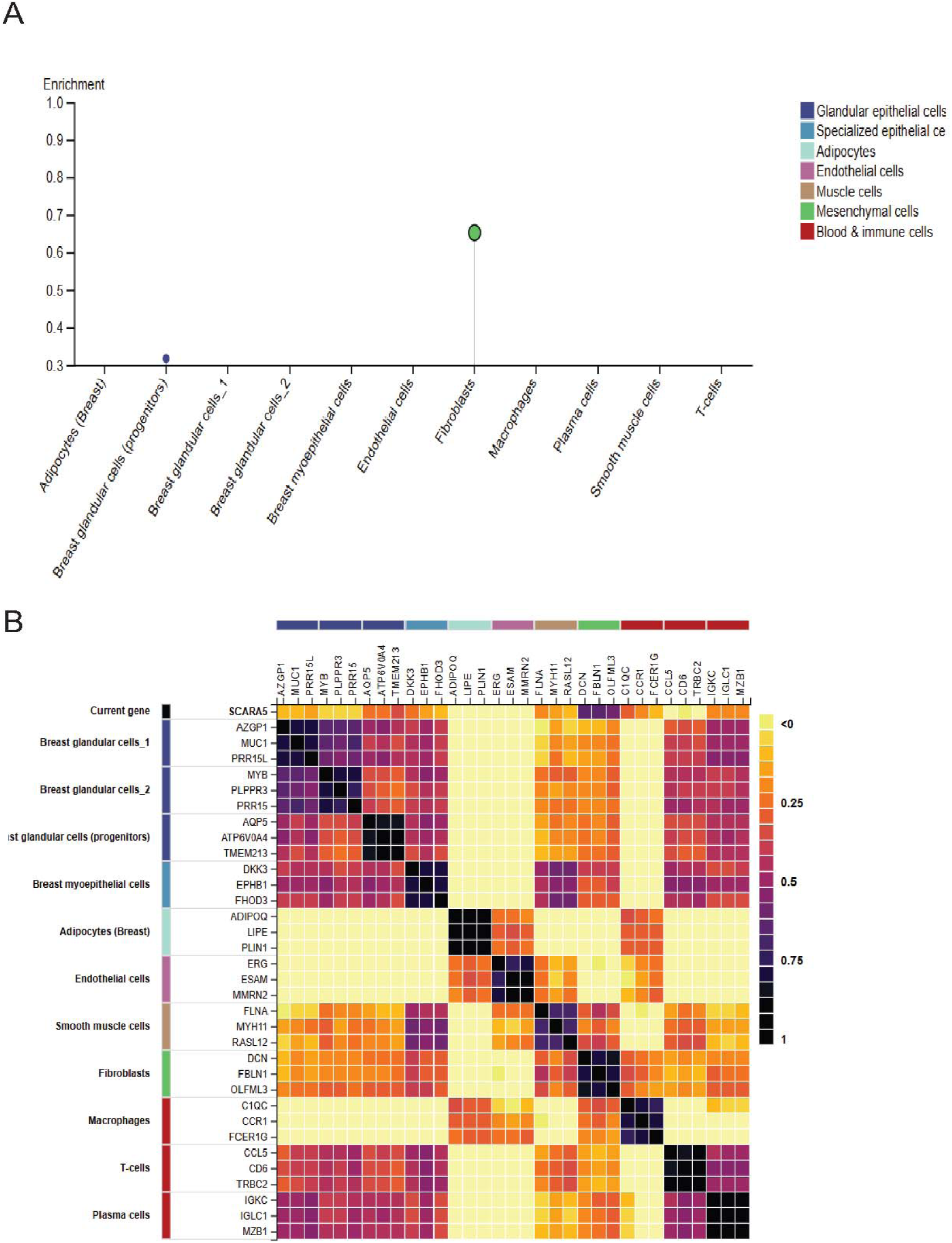
Spatial transcriptomics reveal SCARA5 localization to fibroblast- and endothelial-rich niches in breast tissue. **(A)** Cell-type enrichment analysis from the Human Protein Atlas (HPA) dataset showing the highest SCARA5 expression in fibroblasts and endothelial cells. **(B)** Co-expression heatmap aligning SCARA5 with stromal-associated genes and lipid metabolism markers, confirming its spatial restriction to stromal and vascular compartments in breast tissue.

Gene co-expression heatmaps aligned SCARA5 with a fibroblast- and endothelial-enriched signature, displaying high concordance with extracellular matrix-associated genes and lipid metabolism regulators, while showing limited overlap with epithelial or immune lineage markers (Figure 7B). These findings support the notion of SCARA5 as a stromal marker with a preference for mesenchymal and vascular niches within breast tissue.

Importantly, spatial heterogeneity analysis indicated distinct expression hotspots of SCARA5 within fibroblast-rich stromal zones and endothelial compartments, while immune and epithelial regions exhibited low or negligible expression. This suggests that SCARA5’s function may be spatially restricted to stromal-vascular interfaces, potentially contributing to the regulation of tumor–stroma interactions, extracellular matrix remodeling, and immune cell trafficking in the tumor microenvironment.

Together, these data highlight SCARA5’s spatially restricted expression pattern, reinforcing its proposed role in mediating stromal and vascular dynamics within breast tissue, a feature that may underpin its contribution to tumor suppression and immune-metabolic regulation in the breast cancer milieu.

## 4. DISCUSSION

This study systematically examined the expression, functional connections, and clinical importance of SCARA5 in breast cancer through a multi-omics integrated approach. Our results show a consistent decrease of SCARA5 in breast tumors, especially in aggressive subtypes like HER2-enriched and Luminal B, and highlight its preferred expression within stromal fibroblasts and endothelial cells of the tumor microenvironment (TME). This aligns with SCARA5’s known role as a tumor suppressor in other cancers, including hepatocellular carcinoma, renal cell carcinoma, gastric cancer, oral squamous cell carcinoma, lung cancer, and melanoma (14–16, 21–23, 34, 35).

The tumor microenvironment (TME) plays a crucial role in cancer progression, metastasis, and therapeutic resistance, serving as a dynamic setting where stromal, immune, and tumor cells interact (36–40). Our results indicate that SCARA5 is mainly expressed in stromal fibroblasts and endothelial cells, but not in malignant epithelial cells, implying that its tumor-suppressive function may be mediated through regulation of stromal activities within the TME, as others have suggested (22, 41). This is especially relevant given the growing recognition of cancer-associated fibroblasts (CAFs) and endothelial cells as active participants in tumor growth, immune evasion, and therapy response modulation (42–44).

Functional enrichment and co-expression analyses consistently place SCARA5 within pathways related to lipid metabolism, immune regulation, and scavenger receptor activity. The strong link with lipid metabolism genes such as FABP4, ADIPOQ, and CD36 is significant, as changes in lipid handling in the TME have been connected to metabolic reprogramming that promotes tumor growth and immune suppression (45–47). Additionally, SCARA5’s association with immune-related genes like CLEC3B and LYVE1 indicates it may play a role in shaping the immunomodulatory environment of the TME, a theory supported by the known functions of stromal cells in influencing immune responses (48–52). The functional connection to PPAR signaling and adipocytokine pathways further underscores the complex interaction between metabolism and immunity within the tumor stroma, which could impact disease progression and treatment responses (53–57).

These findings have significant translational implications. The observed downregulation of SCARA5 in aggressive breast cancer subtypes, along with its association with poor patient outcomes, suggests its potential as a prognostic biomarker, especially in HER2-enriched and Luminal B breast cancers, both of which are known for high relapse rates and therapeutic resistance (58–61). Furthermore, the stromal-specific expression pattern of SCARA5 indicates that therapeutic strategies aimed at modulating stromal function—such as targeting CAFs or remodeling the extracellular matrix—might benefit from including SCARA5 status to improve patient selection. The connection of SCARA5 to metabolic and immune pathways also opens the possibility of using existing drugs that target PPAR signaling, lipid metabolism, or immune checkpoints in combination with stroma-modulating approaches (62).

Single-cell and spatial transcriptomics analyses highlighted the compartmentalization of SCARA5 expression within fibroblast- and endothelial-rich niches, with minimal expression detected in epithelial or immune cells. This spatial restriction supports the idea of stromal specialization in regulating extracellular matrix integrity, vascular dynamics, and immune cell trafficking (63). For example, recent primary studies have revealed stromal specialization mechanisms, such as glycolytic CAFs creating CXCL16-mediated ECM barriers that limit CD8 T cell infiltration (64), and Hyaluronan-rich cancer-associated fibroblast (CAF) matrices directing macrophage movement and influencing angiogenesis through ECM restructuring and vascular remodeling (65). The ability of SCARA5 to impact these processes suggests its potential as a therapeutic target, especially in strategies aiming to disrupt tumor–stroma interactions or normalize the TME to improve drug delivery and immune cell infiltration. Incorporating SCARA5 into spatial transcriptomic diagnostics may also assist in patient stratification and monitoring treatment responses within precision oncology frameworks.

Nonetheless, this study has limitations. Although our insilico analyses revealed consistent associations across diverse datasets, functional validation in cellular and in vivo models is essential to confirm SCARA5’s mechanistic role in stromal regulation and its impact on tumor behavior. Moreover, assessing whether modulation of SCARA5 can alter sensitivity to chemotherapy, targeted therapy, or immunotherapy would provide critical insights for its translational application.

## 5. CONCLUSION

Our comprehensive multi-omics investigation positions SCARA5 as a stromal-enriched tumor suppressor with key roles in lipid metabolism, immune regulation, and tumor microenvironment modulation in breast cancer. Its consistent downregulation in aggressive subtypes and association with adverse clinical outcomes underscore its potential as both a prognostic biomarker and a therapeutic target. Further functional and translational research on SCARA5 may pave the way for innovative strategies to modulate the tumor stroma, disrupt pro-tumorigenic metabolic pathways, and improve therapeutic efficacy in breast cancer.

## AUTHOR CONTRIBUTIONS

Conceptualization of the study: T.J, S.M, S.K, H.K and MM. Methodology: T.J, S.M, and S.K. Investigation and visualization: T.J. Formal analysis: T.J. Writing original draft: T.J. Review and editing: T.J, S.M, M.M. Supervision: H.K and M.M.

## DATA AVAILABILITY STATEMENT

All data supporting the findings of this study are available within the paper and its supplementary information file.

## ETHICS APPROVAL STATEMENT

None

## LIST OF SUPPORTING INFORMATION

Table S1: List of top 50 differentially expressed genes (DEGs) identified in breast cancer versus normal breast tissues from TCGA-BRCA dataset.

Table S2: Reactome pathway enrichment analysis of the top 50 DEGs, highlighting key biological pathways associated with SCARA5.

Table S3: Detailed list of genes mapped to the enriched Reactome pathways based on the top 50 DEGs.

Table S4: Combined pathway enrichment results (GO, KEGG, and Reactome) for the top 50 DEGs, summarizing significant biological processes and pathways.

